# Integrating multi-OMICS data through sparse Canonical Correlation Analysis for predicting complex traits: A comparative study

**DOI:** 10.1101/843524

**Authors:** Theodoulos Rodosthenous, Vahid Shahrezaei, Marina Evangelou

## Abstract

**Motivation:** Recent developments in technology have enabled researchers to collect multiple OMICS datasets for the same individuals. The conventional approach for understanding the relationships between the collected datasets and the complex trait of interest would be through the analysis of each OMIC dataset separately from the rest, or to test for associations between the OMICS datasets. In this work we show that by integrating multiple OMICS datasets together, instead of analysing them separately, improves our understanding of their in-between relationships as well as the predictive accuracy for the tested trait. As OMICS datasets are heterogeneous and high-dimensional (*p* >> *n*) integrating them can be done through Sparse Canonical Correlation Analysis (sCCA) that penalises the canonical variables for producing sparse latent variables while achieving maximal correlation between the datasets. Over the last years, a number of approaches for implementing sCCA have been proposed, where they differ on their objective functions, iterative algorithm for obtaining the sparse latent variables and make different assumptions about the original datasets.

**Results:** Through a comparative study we have explored the performance of the conventional CCA proposed by Parkhomenko et al. [2009], penalised matrix decomposition CCA proposed by Witten and Tibshirani [2009] and its extension proposed by Suo et al. [2017]. The aferomentioned methods were modified to allow for different penalty functions. Although sCCA is an unsupervised learning approach for understanding of the in-between relationships, we have twisted the problem as a supervised learning one and investigated how the computed latent variables can be used for predicting complex traits. The approaches were extended to allow for multiple (more than two) datasets where the trait was included as one of the input datasets. Both ways have shown improvement over conventional predictive models that include one or multiple datasets.

## 1 Introduction

Nowadays, it is becoming a common practice to produce multiple OMICS (*e*.*g*. Transcriptomics, Metabolomics, Proteomics etc.) datasets from the same individuals [TCGA, 2012, Inouye et al., 2010, Lenz et al., 2008] leading to research questions involving the in-between relationships of the datasets as well as with the complex traits (responses). The datasets obtained through different mechanisms lead to different data distributions and variation patterns. The statistical challenge is how can these heterogeneous and high-dimensional datasets be analysed to understand their in-between relationships. A follow up question to address is how can these relationships be used for understanding the aetiology of complex traits.

Over the past years a number of data integration approaches have been proposed for finding in-between dataset relationships [Lock et al., 2013, Niu et al., 2017]. These approaches can be characterised by their strategy: (A) Early: Combining data from different sources into a single dataset on which the model is built, (B) Intermediate: Combining data through inference of a joint model, and (C) Late: Building models for each dataset separately and combining them to a unified model [Gligorijević and Pržulj, 2015]. A number of data integration approaches have been proposed in the literature for clustering disease subtypes [Swanson et al., 2019, Mariette and Villa-Vialaneix, 2018]. Huang et al. [2017] present a review of available methods for data integration and argue the need for direct comparisons of these methods for aiding investigators choosing the best approach for the aims of their analysis. Only few approaches have been proposed in the literature for supervised learning, *i*.*e*. for predicting the disease outcome/response. van Vliet et al. [2012] investigated early, intermediate, and late integration approaches by applying nearest mean classifiers for predicting breast cancer outcome. Their findings suggest that multiple data types should be exploited through intermediate or late integration approaches for obtaining better predictions of disease outcome.

The focus of this paper is on Canonical Correlation Analysis (CCA), an intermediate integrative approach proposed by Hotelling [1936]. CCA and its variations have been applied in various disciplines, including personality assessment [Sherry and Henson, 2005], material science [Rickman et al., 2017], photogrammetry [Vestergaard and Nielsen, 2015], cardiology [Jia et al., 2019], single-cell analysis [Butler et al., 2018].

In the case of integrating two datasets CCA produces two new sets of latent variables, called *canonical (variate) pairs*. Suppose that there are two datasets with measurements made on the same samples (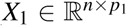 and 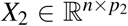, assuming w.l.o.g. *p*_1_ > *p*_2_). For every *i*^*th*^ pair, where *i* = 1, …, min(*p*_1_, *p*_2_), CCA finds two *canonical vectors*, 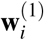 and 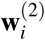, such that 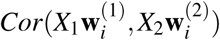 is maximised based on the constraints described below. For the first pair, the only constraint to satisfy is 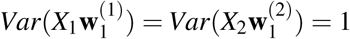. In computing the *r*^*th*^ canonical pair, the following three constraints need to be satisfied:

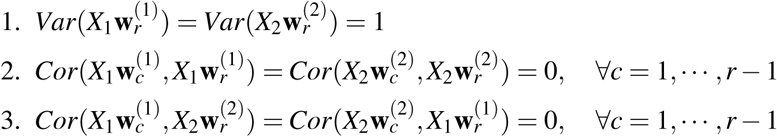

Orthogonality among the canonical variate pairs must hold. That is, not just between the elements of each feature space, but also among all combinations of the canonical variates; except the ones in the same pair, for which the correlation must be maximised. Complete orthogonality is attained when these constraints are satisfied.

In other words, CCA finds linear combinations of *X*_1_ and *X*_2_ that maximise the correlations between the members of each canonical variate pair 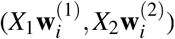, where 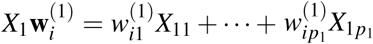 and 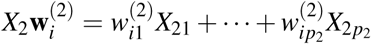. By assuming that there exists some correlation between the datasets, we can look at the most expressive elements of canonical vectors (and indirectly, features) to find relationships between datasets.

CCA can be considered as an extension of Principal Components Analysis (PCA) applied on two datasets rather than one dataset. Similarly to PCA, CCA can be applied for dimension reduction purposes, as the maximum size of the new sets of latent variables is *k* = min{*p*_1_, *p*_2_}.

A solution to CCA can be obtained through Singular Value Decomposition [Hsu et al., 2008]. The canonical vector 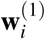 is an eigenvector of 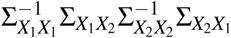, while 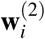 is proportional to 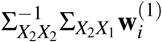. Each pair *i* of canonical vectors corresponds to the respective eigenvalues in a descending order.

In the case of high-dimensional data (*p* >> *n*), the covariance matrix is not invertible and a CCA solution cannot be obtained. Over the years a number of different methods have been proposed in the literature for finding solutions to sCCA. These variations are called sparse CCA (sCCA) methods.

In this paper, three sCCA solutions are discussed, investigated and extended. The first method, Penalised Matrix Decomposition CCA (PMDCCA) proposed by Witten and Tibshirani [2009] obtains sparsity through *l*_1_ penalisation (LASSO Tibshirani [1996]). The bound form of the constraints is used in order to reach a converged solution by iteratively updating **w**^(1)^ and **w**^(2)^. One of the assumptions of the PMDCCA approach is that the two datasets are independent, *i*.*e*. the covariance matrix of each input dataset is assumed to be the identity matrix. The second method proposed by Suo et al. [2017] relaxes the assumption of independence by allowing dependent data to be analysed through proximal operators (for the rest of the manuscript we are referring to this method as RelPMDCCA). Even though, the additional restrictions make RelPMDCCA practically more applicable, it is computationally more expensive than PMDCCA.

The third sCCA method we investigated is Conventional CCA (ConvCCA) proposed by Parkhomenko et al. [2009]. Similarly to PMDCCA sparsity is obtained through LASSO penalisation. ConvCCA estimates the singular vector of 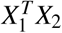 while iteratively applies the soft-thresholding operator. Chalise and Fridley [2012] extended ConvCCA to allow for other penalty functions and through a comparative study found the Smoothly Clipped Absolute Deviation (SCAD) penalty to produce the most accurate results. Motivated by this finding, in our work we have modified both ConvCCA and RelPMDCCA to be penalised through SCAD.

Section 2 starts with a description of the three sCCA approaches: PMDCCA, RelPMDCCA, ConvCCA, followed by a description of our proposed extension of ConvCCA and RelPMDCCA to allow for multiple input datasets. A comprehensive simulation study has been conducted for comparing the performance of the three methods and their extensions on integrating two and multiple datasets. To our knowledge, a comparison of the three sCCA methods has not been made elsewhere. The simulated datasets and scenarios considered are presented in Section 3.1.

We have further addressed the second important question of how can data integration be used for linking the multi-OMICS datasets with complex traits/responses? For addressing this question we have looked the problem as both supervised and unsupervised one. For the supervised model, we have used the computed canonical pairs as input predictors in regression models for predicting the response. Also, we have explored the possibility of adding the response vector as an input matrix in the setting of multiple dataset integration. We have found both these approaches to have a better predictive accuracy than conventional machine learning methods that use either one or both of the input datasets. Section 4 presents the analysis of real datasets with the aim of predicting traits through sCCA.

## 2 Methods

All sCCA methods share a common objective function, given by:

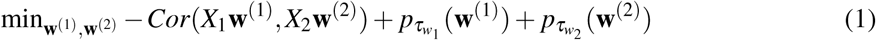

where 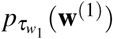 and 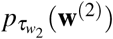 represent penalty functions on **w**^(1)^ and **w**^(2)^ respectively. It is a bi-convex optimization where, if **w**^(1)^ is fixed, then eq. (1) is convex in **w**^(2)^ and vice versa. Hence, one can find a solution through an iterative algorithm.

In this section, the computation of the first canonical pair is presented. To derive additional pairs, we have extended an approach proposed by Suo et al. [2017], which is presented in Section 2.6.

### 2.1 Conventional CCA (ConvCCA)

Parkhomenko et al. [2009] proposed a solution of sCCA based on approximating the sample correlation matrix and applying LASSO penalisation through the soft-thresholding operator proposed by Tibshirani [1996]. An iterative procedure updates both canonical vectors, **w**^(1)^ and **w**^(2)^ at each iteration *k*. The procedure is illustrated in the following steps where one of the vectors (*e*.*g*. **w**^(1)^) is updated while the second (*e*.*g*. **w**^(2)^) is kept fixed:

1. Compute sample correlation matrix 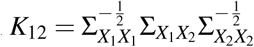
2. (**w**^(1)^)^*k*+1^ ← *K*_12_(**w**^(2)^)^*k*^
3. Normalise 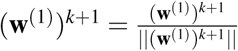
4. Apply soft-thresholding: 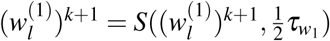
5. Normalise 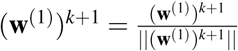

where 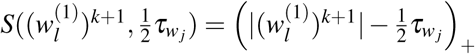*Sign* 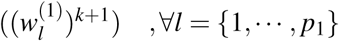 and

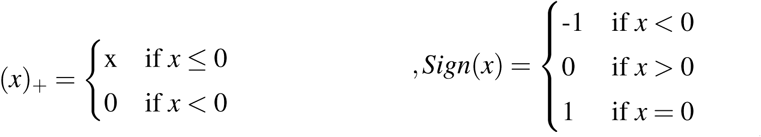

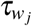 represents the tuning parameter for each dataset *X*_*j*_ (*j* = 1, 2) and (**w**^(1)^)^*k*^ is the value of **w**^(1)^ at the *k*^*th*^ iteration. To update **w**^(2)^, the same procedure is followed with the difference of replacing the second step with 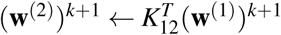.

### 2.2 Penalised Matrix Decomposition CCA (PMDCCA)

Witten and Tibshirani [2009] formulated the problem as:

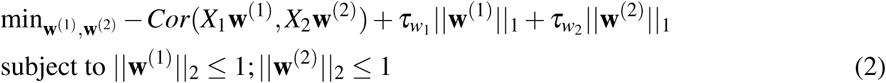

an iterative algorithm based on Penalised Matrix Decomposition (PMD) is applied with the update formula:

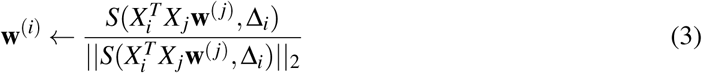

where Δ_*i*_ > 0 is chosen such that ‖**w**^(*i*)^‖_1_ = 1 holds, or Δ_*i*_ = 0 if ‖**w**^(*i*)^‖_1_ ≤ 1, for *i, j* = 1, 2, *i* ≠ *j*. Although the PMDCCA solution is different from the ConvCCA solution both approaches assume that the features are independent within each dataset (*i*.*e*. 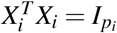, *i* = 1, 2).

### 2.3 Relaxed PMDCCA (RelPMDCCA)

Suo et al. [2017] proposed a solution that relaxes the independence assumption and applies penalisation through proximal operators. Their formulation of the problem is similar to (2):

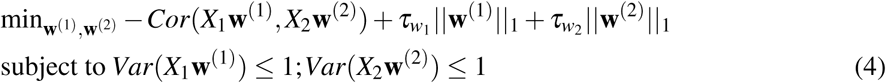

The solution to this optimisation is obtained through linearised Alternating Direction Method of Multipliers [Parikh, 2014, Boyd et al., 2010].

The iterative updates on the canonical variate pairs by RelPMDCCA are given as follows:

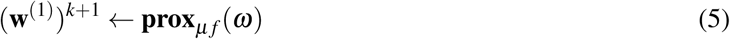

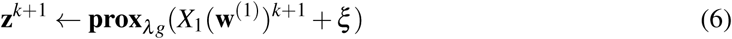

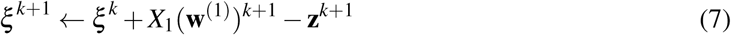

where:

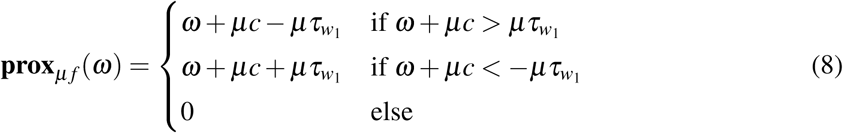

and

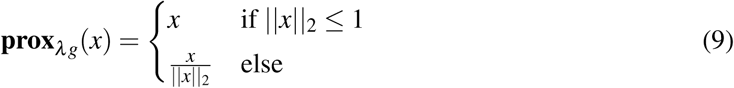

with 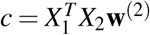 and 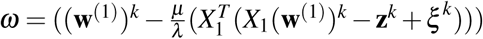. The parameter 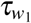 controls the sparseness level while the algorithm parameters *µ* and *λ* must satisfy 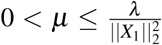 [Parikh, 2014]. **w**^(2)^ is updated through the same proximal operators, with 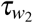 acting as the tuning parameter.

### 2.4 Implementing SCAD penalty

The Smoothly Clipped Absolute Deviation (SCAD) [Fan and Li, 2001] penalty with tuning parameter *τ* applied on *w* is given as follows:

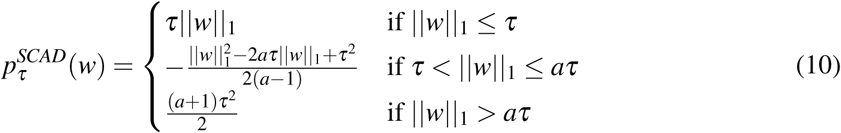

where *a* is fixed and suggested by Fan and Li to be set as *a* = 3.7. Motivated by the findings of Chalise and Fridley (2013), we have modified RelPMDCCA to perform penalisation through SCAD.

In the objective functions of ConvCCA (eq. 1) and RelPMDCCA (eq. (4)), 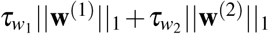 is replaced by 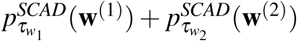. As a result the iterative updates of the canonical vectors are different. In ConvCCA, the algorithm is adjusted accordingly by replacing the soft-thresholding operator in step 4 with:

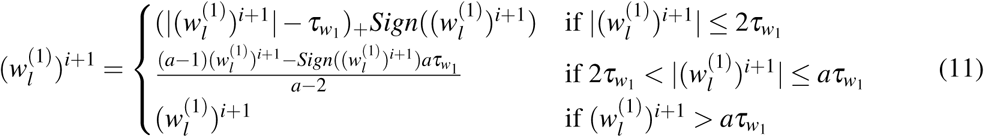

The updates of canonical vectors in RelPMDCCA with SCAD penalty are performed by:

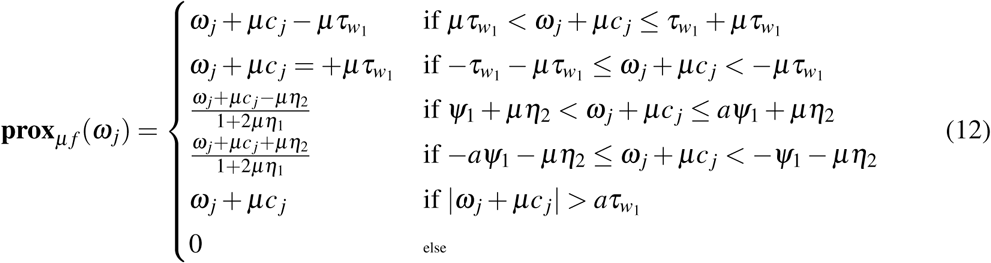

where 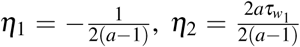,and 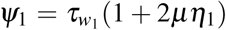. The update functions of **z** and *ξ* remain the same.

### 2.5 Multiple sCCA

In OMICS studies, it common for a study to have more than two datasets (such as transcriptomics, genomics, proteomics and metabolomics) on the same patients. Incorporating all available data simultaneously through an integrative approach can reveal unknown relationships between the datasets. This section presents extensions of the three sCCA methods we have seen, for the integration of multiple (more than two) datasets simultaneously.

Suppose we have *M* separate datasets denoted by *X*_1_, …, *X*_*M*_, where 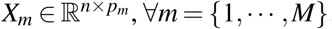. The problem of eq. (1) is then generalised as:

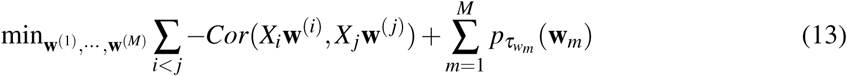

As sCCA is bi-convex, multiple sCCA is multi-convex, *i*.*e*. if **w**^(*j*)^, ∀ *j* ≠ *i* are fixed, then the problem is convex in **w**^(*i*)^. Instead of producing maximal correlated canonical variate pairs (2-tuple), multiple sCCA produces canonical variate list (*M-tuple*), *e*.*g*., for *i* = 1, …, min(*p*_1_, …, *p*_*M*_), the *i*^*th*^ canonical variate list would be 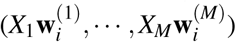. Each *X*_*m*_**w**^(*m*)^ is taken such that, it is maximally correlated with the rest of the latent features Σ_*j*≠*m*_ *X*_*j*_**w**^(*j*)^.

In multiple ConvCCA, we propose to update **w**^(*i*)^ iteratively, by keeping **w**^(*j*)^, ∀ *j* ≠ *i* fixed, as shown in Algorithm 1.

Witten and Tibshirani [2009] proposed an extension to their solution, by assuming that 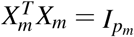, ∀*m* = {1, …, *M*}. To update **w**^(*i*)^, the canonical vectors **w**^(*j*)^, ∀ *j* ≠ *i* are kept fixed. Multiple PMDCCA can then be performed by minimising 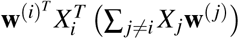, ∀*i* = {1, …, *M*}, with constraint functions ‖**w**^(*m*)^‖_2_ ≤ 1.

We have extended RelPMDCCA for multiple datasets by following the approach of PMDCCA. The constraint functions *Var*(*X*_*m*_**w**^(*m*)^) ≤ 1, ∀*m* = (1, …, *M*) and the proximal operators remain the same. If **w**^(*j*)^ ∀ *j* ≠ 1 are kept fixed, we can update **w**^(1)^, by replacing 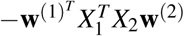 in eq. (4) with 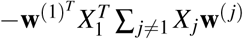.

#### Algorithm 1: Multiple ConvCCA

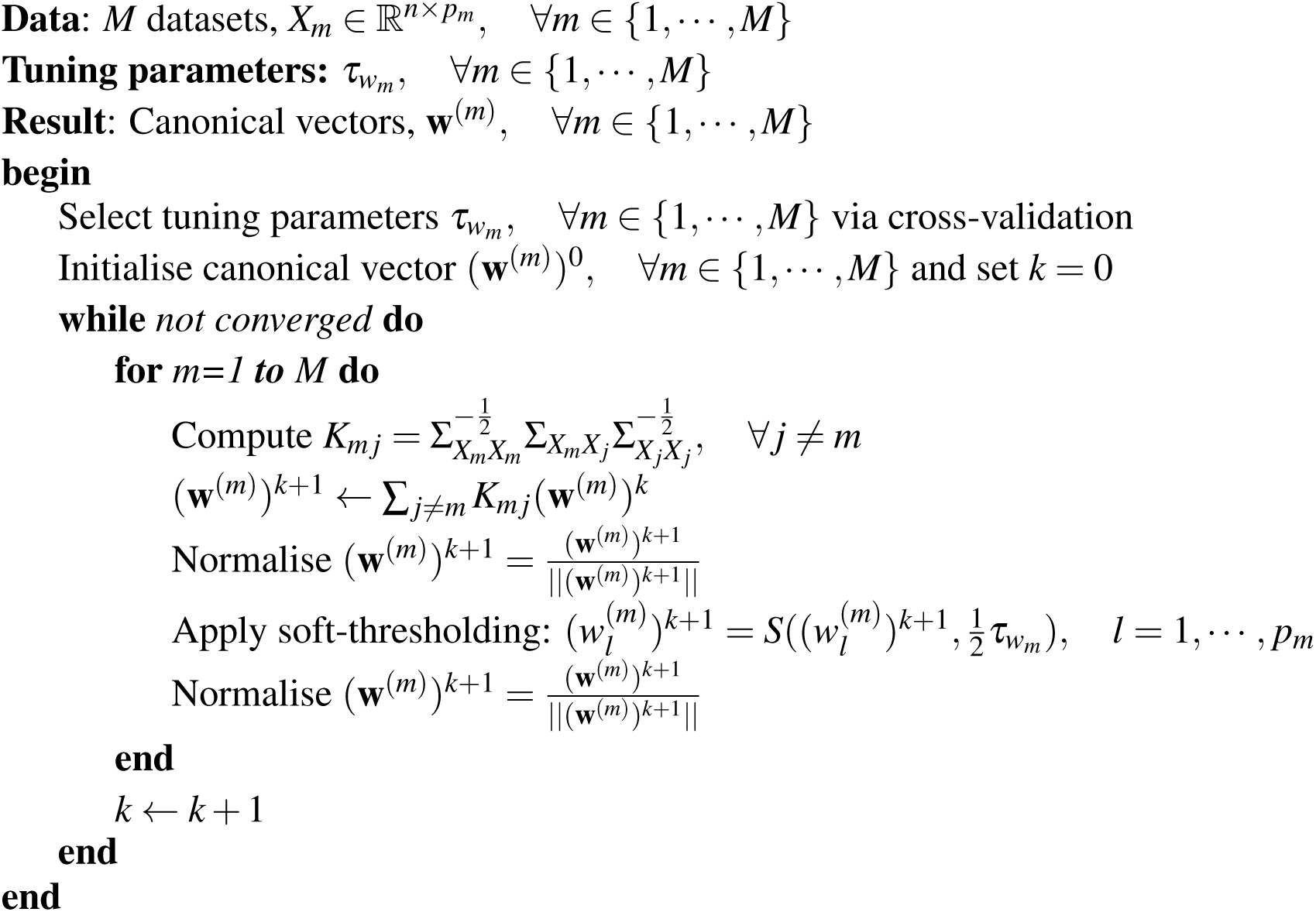

The tuning parameters in sCCA and multiple sCCA are selected as the ones producing maximal correlation through cross-validation. A detailed description of the procedure is presented in the Supplementary material.

In this work we have explored the performance of the sCCA methods for predicting the response of interest by including the response matrix as one of the datasets being integrated.

### 2.6 Computing the additional canonical vectors

We have only addressed the computation of the first canonical variate pair so far but this might not be adequate for capturing the variability of the datasets and of their relationships. Similarly to PCA as the number of computed principal components is increased the total amount of variability explained is increased. By computing the additional canonical vectors of CCA additional constraints must be satisfied as illustrated below.

Suo et al. [2017] compute the remaining canonical vectors by adding the second constraint to the optimisation. Let 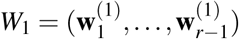 and 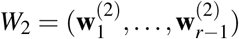 define the *r* − 1 canonical vectors which were computed. The *r*^*th*^ canonical vector is found through the optimisation problem:

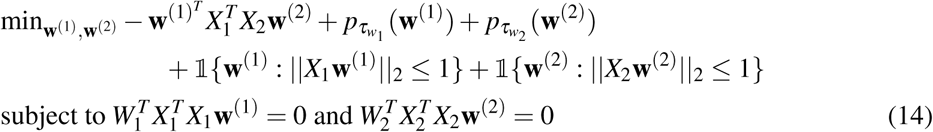

where 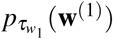 and 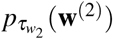 represent the penalty functions on **w**^(1)^ and **w**^(2)^ with parameters 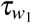 and 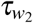 respectively. This solution would successfully result in producing latent features that are uncorrelated within the new datasets of canonical vectors, although the correlation is not restricted between the two new datasets.

In an attempt to include the additional constraint to the optimisation problem, we propose the following extension to eq. 14:

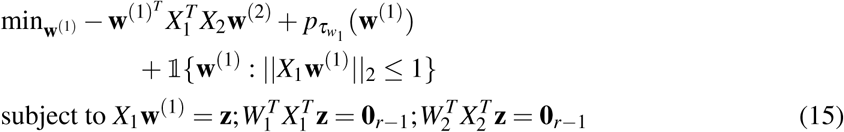

The solution to this optimization problem is presented in Algorithm 2.

#### Algorithm 2: Computing the additional canonical vectors

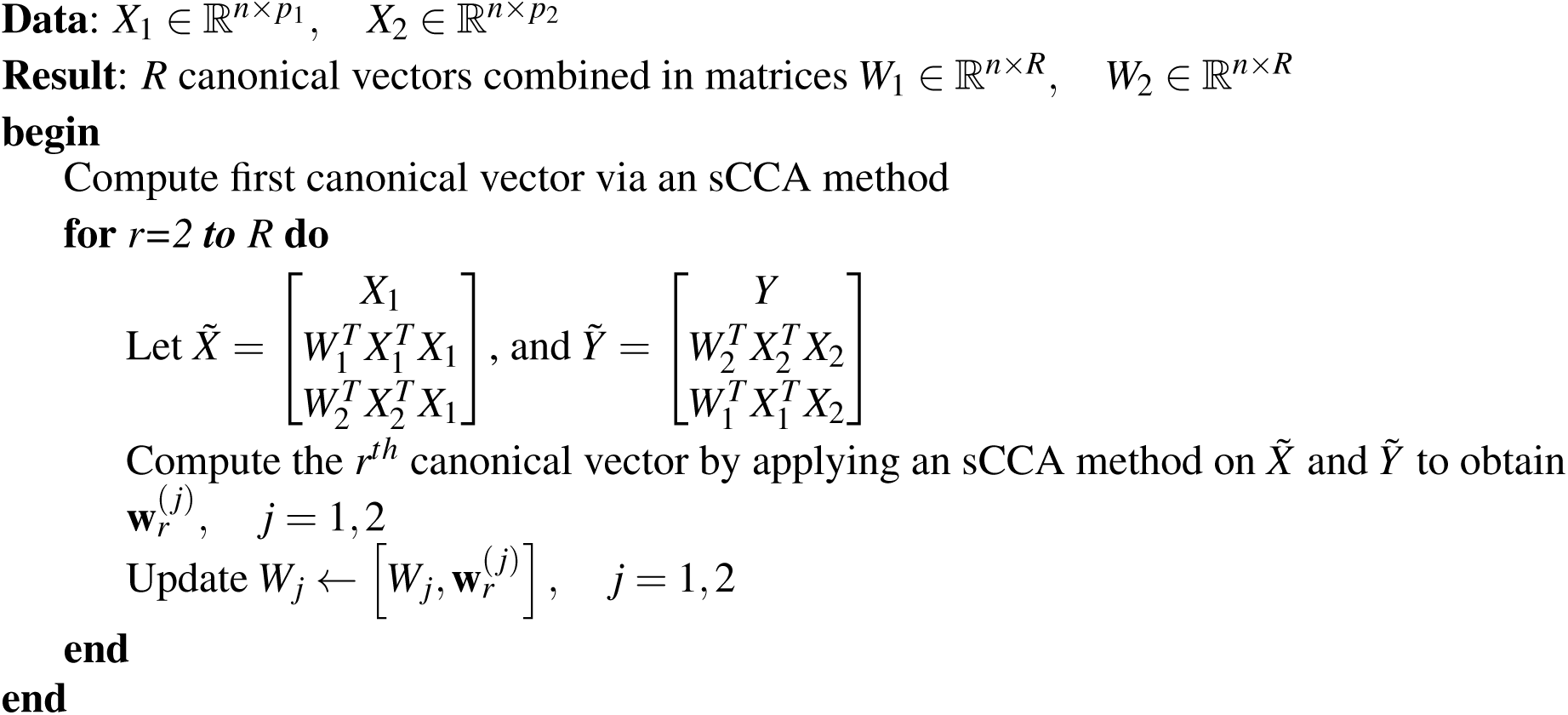

Witten and Tibshirani [2009] proposed to update the cross-product matrix 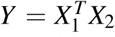 after the computation of each canonical pair, by 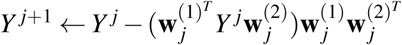, where 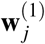 and 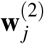 are the *j*^*th*^ canonical vectors. The authors of ConvCCA proposed to take the residual of *K* by removing the effects of the first canonical vectors and repeat the algorithm in order to obtain the additional canonical pairs.

## 3 Simulation study

Simulated datasets were generated for assessing the performance of the three methods on: (i) integrating two datasets, (ii) the orthogonality attained by each method and (iii) integrating multiple datasets.

### 3.1 Models, Scenarios & Evaluation Measures

#### 3.1.1 Models

Three models were used for simulating data with similar characteristics as OMICS datasets. Different types of scenarios were examined covering a range of possible data characteristics. All three data generating models are based on five parameters, 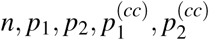, where *n* represents the number of samples, *p*_*i*_ is the total number of features in *X*_*i*_ and 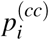 represents the number of features in *X*_*i*_ which are cross-correlated with the rest of datasets 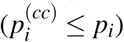.

##### 3.1.1.1 Simple Model

A simple data generating model that generates data for *M* ≥ 2 datasets:

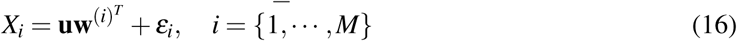

where **u** ∈ ℝ^*n*^, 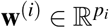 and *ε*_*i j*_ ∼ *𝒩* (0, 1), *j* = 1, …, *p*_*i*_. Only the first 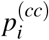 elements of **w**^(*i*)^ are non-zero, representing the cross-correlated features that we seek to identify.

##### 3.1.1.2 Single-Latent Variable Model

Parkhomenko et al. [2009] proposed a single-latent variable model in assessing ConvCCA. An extension of this model is presented here, allowing the generation of multiple datasets (see Figure 1). *M* datasets, 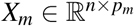, *m* = {1, …, *M*} are generated, such that the first 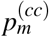 features of each *X*_*m*_ would be cross-correlated. In other words, w.l.o.g. the first 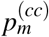 of *X*_*m*_ will be correlated with the first 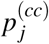 of *X*_*j*_, ∀ *j* ≠ *m*. These groups of features are associated with each other according to the same (single-latent variable) model. A latent variable *w*^(*i*)^, explains a subset of observed variables in *X*_*i*_, i.e. 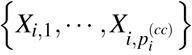. Through a common higher level latent variable, *µ, w*^(*i*)^ ∀*i*, are correlated. The rest of the features are independent within their respective datasets.

**Figure 1:**
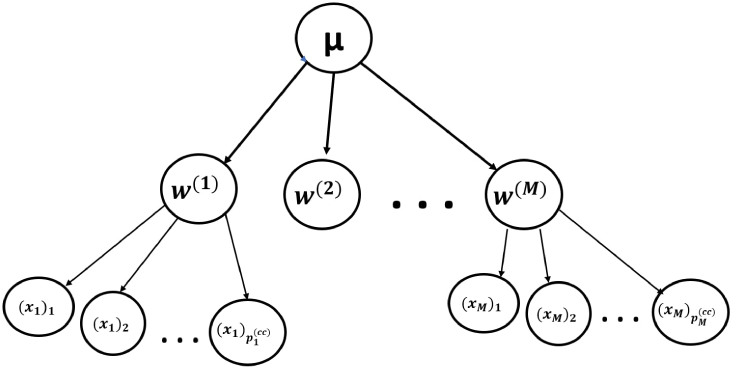
The single latent variable model simulation

After simulating a random variable 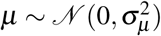, the data are generated as follows:

1. For the cross-correlated variables:

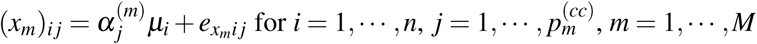

where we assume 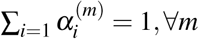, and 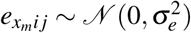, ∀*i, j, m*.
2. For the independent variables:

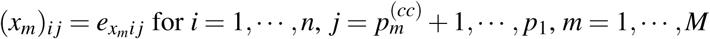

where again we assume 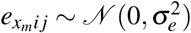, ∀*i, j, m*.

##### 3.1.1.3 Covariance-based Model

Suo et al. [2017] proposed simulations based on the structure of the covariance matrices of both datasets (*X*_1_ and *X*_2_). Three types of covariance matrices were considered in this study: (i) Identity, (ii) Toeplitz and (iii) Sparse. We have utilised this model for generating two datasets. Suppose, 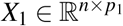 and 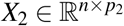 with data being generated by:

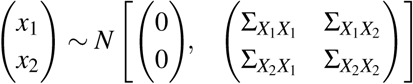

where 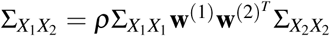, with **w**^(1)^ and **w**^(2)^ being the true canonical vectors and *ρ* the true canonical correlation. The covariance matrices 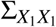 and 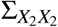 are explicitly defined based on the type of data generating model.

#### 3.1.2 Scenarios and Evaluation Measures

In comparing the three sCCA methods for integrating two datasets, six scenarios of different data characteristics were examined (Table 1). The first scenario acts as a baseline for the rest. A single parameter is changed for each additional scenario.

**Table 1:** Data characteristics and simulation scenarios used to evaluate the three sCCA methods for integrating two datasets.

| Simulation Scenarios |  |  |  |  |  |  |
| --- | --- | --- | --- | --- | --- | --- |
| Scenarios | Data Characteristics |  |  |  |  | Parameters |
| 1 | $n = 40,$ | $p_1 = 80,$ | $p_2 = 60,$ | $p_1^{(cc)} = 5,$ | $p_2^{(cc)} = 15$ | $n$: The number of samples |
| 2 | $n = \mathbf{150},$ | $p_1 = 80,$ | $p_2 = 60,$ | $p_1^{(cc)} = 5,$ | $p_2^{(cc)} = 15$ | $p_i$: The number of features in |
| 3 | $n = 40,$ | $p_1 = \mathbf{200},$ | $p_2 = 60,$ | $p_1^{(cc)} = 5,$ | $p_2^{(cc)} = 15$ | dataset $i$, for $i = 1, 2$ |
| 4 | $n = 40,$ | $p_1 = 80,$ | $p_2 = \mathbf{200},$ | $p_1^{(cc)} = 5,$ | $p_2^{(cc)} = 15$ | $p_i^{(cc)}$: The number of |
| 5 | $n = 40,$ | $p_1 = 80,$ | $p_2 = 60,$ | $p_1^{(cc)} = \mathbf{50},$ | $p_2^{(cc)} = 15$ | cross-correlated features |
| 6 | $n = 40,$ | $p_1 = 80,$ | $p_2 = 60,$ | $p_1^{(cc)} = 5,$ | $p_2^{(cc)} = \mathbf{50}$ | in dataset $i$, for $i = 1, 2$ |

We assessed the performance of the sCCA methods for integrating multiple datasets by generating three datasets through three scenarios: (i) *n* = 40, *p*_1_ = 80, *p*_2_ = 60, *p*_3_ = 40, 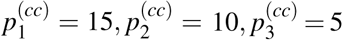, (ii) *n* = 40, **p**_1_ **= 200**, *p*_2_ = 60, *p*_3_ = 40, 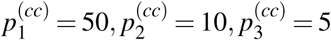, (iii) **n = 150**, *p*_1_ = 80, *p*_2_ = 60, *p*_3_ = 40,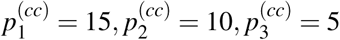.

The sCCA methods in both simulation studies were evaluated by measuring: (i) the canonical correlation, (ii) the correct identification of sparsity in the data, by computing accuracy, precision and recall of the true non-zero elements of the estimated canonical vectors, and (iii) the loss between true and estimated canonical vectors. A detailed description of the evaluation measures is presented in the Supplementary material.

An additional simulation study was conducted to evaluate orthogonality. Two datasets were generated, with each of the following scenarios used in all three data generating models: (i) *n* = 500, *p*_1_ = 100, *p*_2_ = 200, 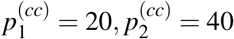, (ii) *n* = 150, *p*_1_ = 100, *p*_2_ = 200, 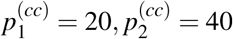 and (iii) *n* = 50, *p*_1_ = 100, *p*_2_ = 200,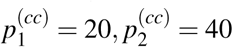.

### 3.2 Results

#### 3.2.1 Integrating two datasets

In the conducted simulation study, the performance of sCCA methods was assessed on the three data generating models and six scenarios shown in Table 1. The results are based on the first canonical pair.

Figure 2a depicts the resulting ROC curves and their Area Under the Curve (AUC) values, averaged over all data generating models and scenarios. ConvCCA with SCAD had the best performance in identifying correctly the sparseness and the non-zero elements of canonical vectors, as it produced the highest AUC. RelPMDCCA with SCAD obtained the lowest AUC value, which shows that the optimal choice of penalty function depends on the sCCA method. While the second dataset (and latent features *X*_2_**w**^(2)^) obtained slightly reduced sensitivity, the sCCA methods performed in a similar fashion as with the first dataset.

**Figure 2:**
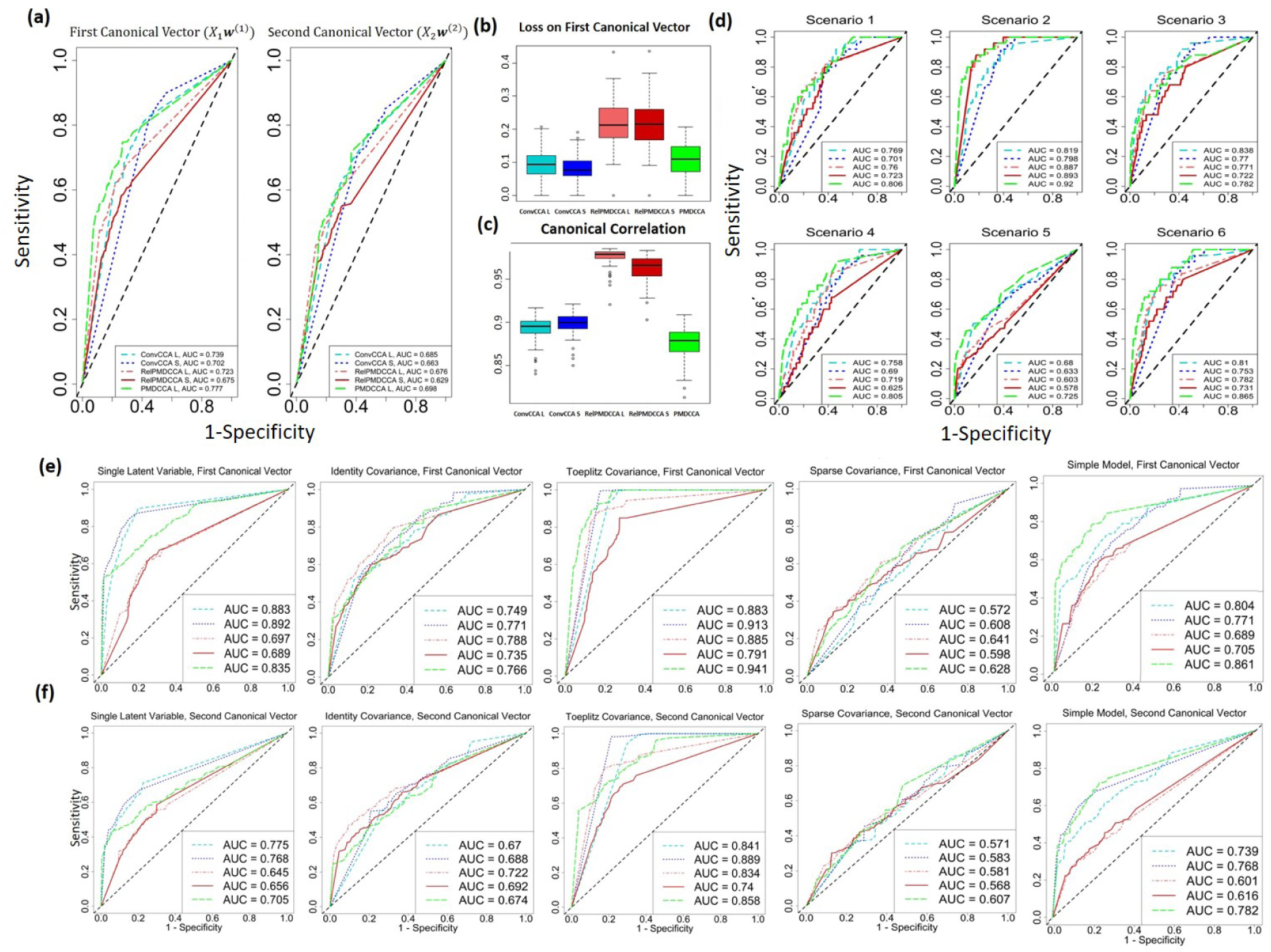
sCCA performance on simulated data for integrating two datasets. **(a)** ROC curve plots on all five sCCA methods after averaging over all data generating models and all scenarios. **(b)** Box-plots of the overall loss of the first canonical vector (*X*_1_**w**^(1)^) averaged over all data generating models and scenarios, and **(c)** canonical correlation in the simulation studies for sCCA. **(d)** ROC curve plots, showing averaged results (over the models) for each scenario on *X*_1_**w**^(1)^. (Results on *X*_2_**w**^(2)^ can be seen in Supplementary material). **(e-f)** ROC curve plots, showing averaged (over the scenarios) results for each model on *X*_1_**w**^(1)^ and *X*_2_**w**^(2)^ respectively.

RelPMDCCA produced the highest loss between the true and estimated canonical vectors (see Figure 2b), but it also provided the highest canonical correlation (Figure 2c). Its overall averaged correlation is close to 1, while for the other two methods, it’s closer to 0.9.

Figure 2d shows the performance of the first canonical vector (*X*_1_**w**^(1)^) averaged over all data generating models. As expected, by increasing the number of samples in a case where *n* > *p* (Scenario 2), the AUC values on all sCCA increased, with RelPMDCCA showing the largest improvement. However, a decrease in the performance of all sCCA methods is observed when the number of non-zero elements is increased. This can be seen on scenarios 5 and 6, for **w**^(1)^ and **w**^(2)^, respectively. We can argue that in cases where the non-zero elements of a canonical vector are at least half of its length (total number of elements), the methods fail to correctly identify some of them. That might be due to the fact that sCCA methods force penalisation and expect a sparser outcome. Furthermore, since the performance on **w**^(2)^ in scenario 6 is worse than that on **w**^(1)^ in scenario 5, we can argue that the higher the ratio of non-zero elements over the total number of elements, the less accurate the identification. On scenarios 3 and 4, where the total number of features is increased while the non-zero elements are not, sCCA methods performed as well as on the baseline scenario.

After averaging over the scenarios, the methods’ performance on each data generating model is shown in Figures 2e and 2f. The methods’ performance seems to be overall influenced by the choice of data generating model. In the case of single-latent variable model, ConvCCA clearly produced the least errors, while in the simple model, PMDCCA produced the highest AUC values. In the covariance-based models, all sCCA methods performed equally well in estimating correctly non-zero elements of the canonical vectors. Methods on Toeplitz model produced higher AUC values than on Identity model, where Sparse model contained the most errors out of all data generating models. Based on this observation, we can argue that the sparser the data, the less accurate the methods.

#### 3.2.2 Orthogonality and Sparsity

A third simulation study was conducted, with the aim of evaluating orthogonality of the three sCCA methods. The scenarios are split into three cases, based on the data sparsity: (i) *n* > *p*_2_ > *p*_1_, (ii) *p*_2_ > *n* > *p*_1_ and (iii) *p*_2_ > *p*_1_ > *n*.

Table 2 shows the classification of each case with one out of three classes: (A) Full (Orthogonality): All pairs were found to be orthogonal, (B) Partial: Some pairs were found to be orthogonal, (C) None: None of the pairs were found to be orthogonal. Different colours in table 2 refer to the different simulation models: Simple model, Single-latent variable model, Identity Covariance-based model.

**Table 2:** Orthogonality of sCCA methods. The table shows whether the algorithms to obtain the remaining canonical variates performs as it should, *i*.*e*. succeeding in getting orthogonal pairs. **None** refers to not obtaining orthogonality at all, **Full**, refers to obtaining orthogonality between all pairs, **Partial** for some, but not all. Brown represents the simple simulation model, with orange the Single-latent variable model and with blue the Covariance-based model.

| Orthogonality of All Simulations with 5 canonical variates |  |  |  |
| --- | --- | --- | --- |
| Methods – > | PMDCCA | ConvCCA | RelPMDCCA |
| $n > p_2 > p_1$ | None Orthogonality | Full Orthogonality | Partial |
|  | Partial Orthogonality | Partial | None |
|  | Partial | Full | None |
| $p_2 > n > p_1$ | None | None | Full |
|  | Partial | None | Full |
|  | Full | None | Partial |
| $p_2 > p_1 > n$ | Full | None | Full |
|  | Full | None | Full |
|  | Full | None | Full |

**Table 3:** A summary on the performance of sCCA methods based on both the simulation studies conducted and the analysis of real data. The summarised results are split into having two datasets or multiple. It is an intuitive evaluation of the methods, based on the results found in this study.

|  |  |  |
| --- | --- | --- |
| On two datasets | ConvCCA | Great performance on simulation studies, especially on single-latent model.<br>Over-fitted cancerTypes data and performed well on nutriMouse. |
|  | PMDCCA | Good performance on simulation studies, especially on Simple model.<br>Over-fitted cancerTypes data and performed well on nutriMouse. |
|  | RelPMDCCA | Moderate to good performance on simulation studies.<br>Had the best performance in analysing two real datasets. |
| Multiple datasets | ConvCCA | Good performance on simulation studies<br>Avoided over-fitting and improved performance in both data studies. |
|  | PMDCCA | Very good performance on simulation studies.<br>Avoided over-fitting and improved performance in both data studies. |
|  | RelPMDCCA | Moderate to good performance on simulation studies.<br>Overall obtained the best results in both data studies. |

As summarised in Table 2, orthogonality is not always preserved, and that depends on the sCCA method, as well as on the data characteristics. The choice of data generating model did not have a high impact in attaining orthogonality. All sCCA methods were penalised through LASSO during this simulation study.

In case (i), ConvCCA has attained full orthogonality on the first five canonical variate pairs, where PMDCCA and RelPMDCCA failed to do so (except between some pairs). In the other two cases, none of the canonical variates obtained from ConvCCA were orthogonal. PMDCCA preserved complete orthogonality in *p*_2_ > *p*_1_ > *n* case and most when *p*_2_ > *n* > *p*_1_. Complete orthogonality was attained in both of those cases when RelPMDCCA was implemented.

#### 3.2.3 Integrating three datasets

A simulation study on multiple sCCA was performed by generating three datasets through the (i) Simple model and (ii) Single-latent variable model. The same evaluating measures as with the case with the two datasets were computed. Canonical correlation was evaluated by computing the average canonical correlation of each dataset with the rest.

Figure 3a presents the averaged (arithmetic) canonical correlation observed by each method. RelPMDCCA produced the highest, as it did in with the case of two datasets. PMDCCA produced the least correlated and least sparse solution, suggesting that it is not performing very well with multiple datasets where the objective of sCCA is to maximize the correlation between the datasets. Figure 3b shows the sparsity obtained by each method with RelPMDCCA providing the most sparse solution.

**Figure 3:**
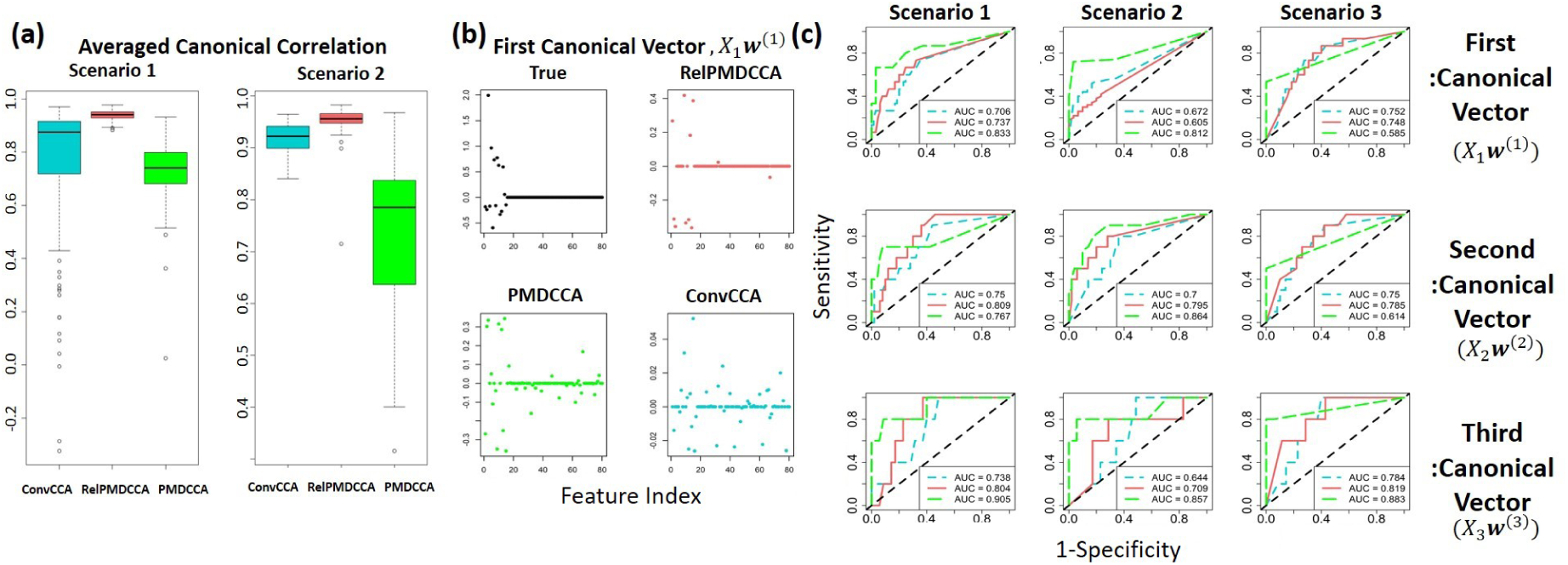
Multiple sCCA performance on simulated data for integrating three datasets. **(a)** Box-plots showing the canonical correlation along the ConvCCA, RelPMDCCA and PMDCCA methods in a multiple-setting. **(b)** An example of a scatter-plot for the first estimated canonical vector. **(c)** ROC curves on multiple sCCA simulations.

Overall, PMDCCA produces the highest AUC values. RelPMDCCA was superior only when the number of samples was increased (Scenario 3). RelPMDCCA and ConvCCA showed a decrease in their performance when the number of non-zero elements was increased, but PMDCCA was able to maintain its good performance.

## 4 Real datasets

### 4.1 NutriMouse

Martin et al. [2007] have performed a nutrigenomic study, with gene expression 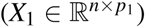 and lipid measurements 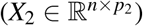 on *n* = 40 mice, with *p*_1_ = 120 genes and concentrations of *p*_2_ = 21 lipids were measured. Two response variables are available: diet and genotype of mice. Diet is a 5-level factor: *coc, fish, lin, ref, sun* and genotype is recorded as either *Wild-type* (WT), or *Peroxisome Proliferator-Activated Receptor-α* (PPAR*α*). NutriMouse data are perfectly balanced in both responses, in that an equal number of samples is available for each class.

Through the analysis of the NutriMouse datasets we aimed to: (1) evaluate which of three considered sCCA approaches performs better, and (2) determine whether data integration of datasets can improve prediction over conventional approaches that only analyse a single dataset. For addressing the second question we have implemented the following off the shelf statistical machine learning approaches: (A) Principal Components Regression (PCR) - logistic model with first 10 principal components acting as predictors, (B) Sparse Regression (SpReg) - penalised (through LASSO and SCAD) logistic and multinomial models, when the response was diet and genotype respectively, (C) k-Nearest Neighbours (kNN) for supervised classification and (D) k-means for unsupervised clustering - acting as a benchmark in splitting the data by ignoring the labels. Since two datasets are available, these four methods were implemented on the following three cases of input data: (i) only *X*_1_, (ii) only *X*_2_, (iii) 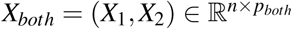, where *p*_*both*_ = *p*_1_ + *p*_2_.The aforementioned machine learning methods and the three sCCA approaches were applied on 100 bootstrap samples of the datasets, taking separate training and test sets at each repetition, for assessing the predictive accuracy of the methods. For the machine learning method applied, the version with the input dataset obtaining the smallest error was compared with the sCCA approaches. In predicting the genotype response using only *X*_1_ (*i*.*e*. gene expression data) was preferred, but *X*_*both*_ was chosen with diet acting as the response (see Supplementary material).

Figure 4 shows the predictive accuracy of the methods on both diet (Figure 4a) and genotype (Figure 4b). PCR and k-means were the least accurate methods (see Supplementary material). In predicting either response, sCCA methods outperformed the conventional machine learning methods. Both the sCCA approaches and the conventional machine learning approaches had very high accuracy for predicting the genotype response whereas their accuracy was lower for predicting the diet response. All the methods had an accuracy between 0.7 and 0.86, except RelPMDCCA that had the highest accuracy 0.92. The precision and recall measures showed similar patterns with RelPMDCCA obtaining the highest values for both measures against all other methods (see Supplementary Material).

**Figure 4:**
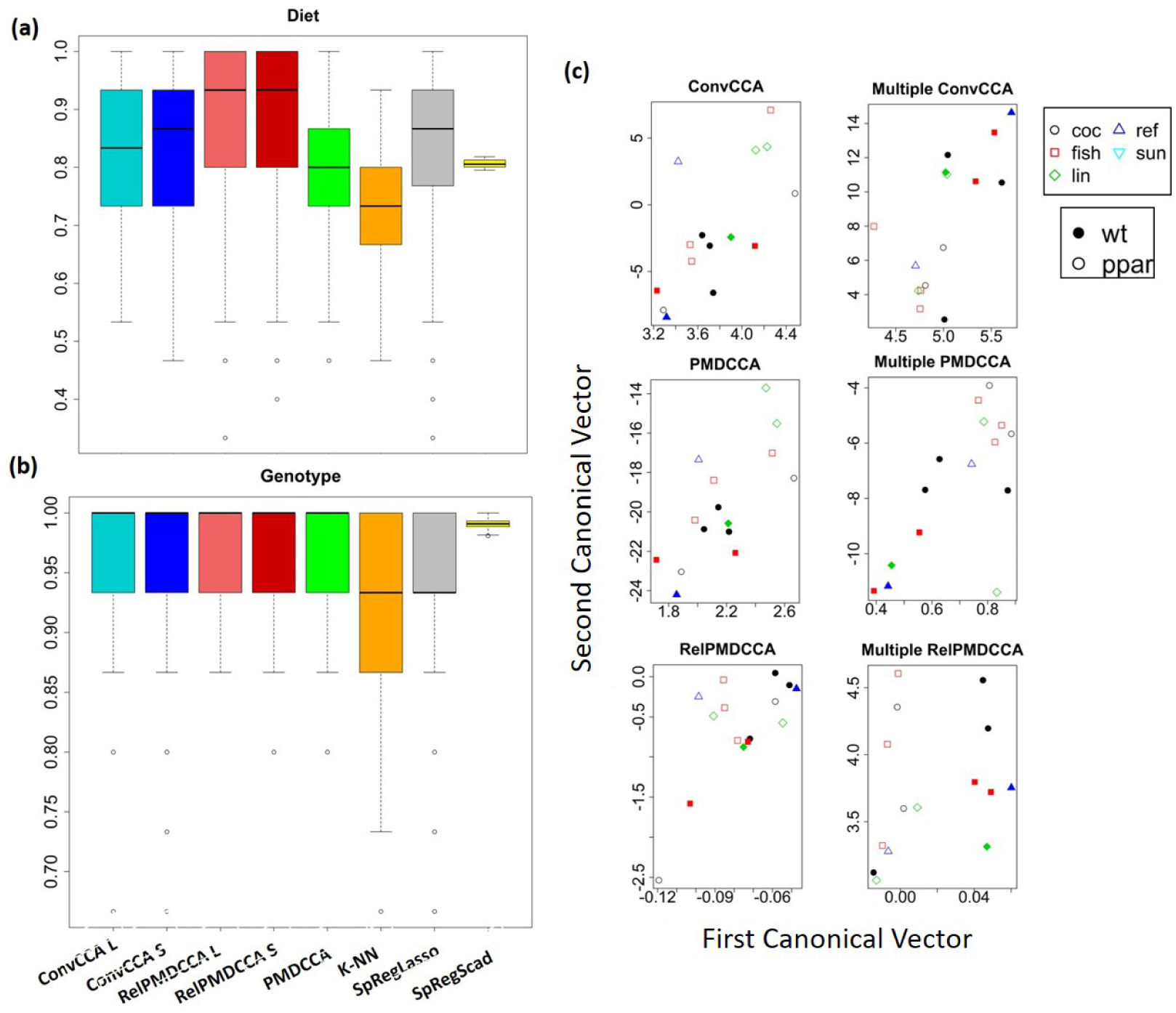
sCCA performance on nutriMouse data. **(a-b)** Box-plots presenting the accuracy of sCCA methods, k-NN, and SpReg with LASSO and SCAD with the response being **(a)** diet and **(b)** genotype. **(c)** Scatter plots of the canonical vectors from the first canonical variate pair of a random nutriMouse test set, after applying sCCA and multiple sCCA.

#### Multiple sCCA with response matrix

We applied our proposed extensions of the sCCA approaches for multiple datasets, where one of the input datasets is the matrix of the two response vectors.

Figure 4c presents the canonical vectors of the first canonical variate pair. The first column of plots shows the canonical vectors obtained without considering a response matrix. On a two-setting integration, the data are not separated well for neither response. However, by including the response matrix, multiple sCCA methods separated clearly the samples between WT and PPAR*α* mice, as shown in the second column of Figure 4c. A slight improvement in their separation between diets was also observed. All multiple sCCA methods performed equally well, although visually, RelPMDCCA indicates the clearest separation in terms of genotype.

### 4.2 CancerTypes

Due to the abundance of data in The Cancer Genome Atlas (TCGA) database, a lot of researchers have applied various integrative algorithms for cancer research [Parimbelli et al., 2018, Wang et al., 2014, Lock et al., 2013, Poirion et al., 2018]. Gene expression, miRNA and methylation data from three separate cancer types were taken: (i) Breast, (ii) Kidney, and (iii) Lung. For each patient, we also have information about their survival status. Thus, the goal of this analysis was to assess whether (multiple) sCCA can improve conventional classification methods in determining the cancer type, and survival status.

The data consist of 65 patients with breast cancer, 82 with kidney cancer and 106 with lung cancer, from which 155 patients are controls. The data in this study covers 10299 genes (*X*_1_), 22503 methylation sites (*X*_2_) and 302 mi-RNA sequences (*X*_3_). Data cleaning techniques, such as removing features with low gene expression and variance were used, leaving us with a remaining of 2250 genes and 5164 methylation sites. Similarly to Section 4.1, k-NN, PCR and SpReg were applied, along with sCCA algorithms. After investigating the best combination, miRNA expression and methylation datasets were selected for integrating two datasets. Multiple sCCA was implemented on all three available datasets.

Table 5a presents the accuracy, precision and recall of each method in predicting the patient’s survival status. The results for predicting cancer type are presented in the Supplementary Material. SpReg, ConvCCA and PMDCCA produced perfect recall, while their precision was recorded around 60%. Such finding suggests over-fitting as a single class is favoured. Thus, a solution providing good results while avoiding over-fitting would be preferable.

PCR and k-NN did not show any signs of over-fitting, but did not perform well (PCR had accuracy below 0.5 and k-NN had consistently low values on all three measures). RelPMDCCA did not over-fit the data and produced high measure values, especially with LASSO being the penalty function. Multiple RelPMDCCA produced the most accurate and precise solution out of all methods applied. Multiple PMDCCA and ConvCCA improved the results of their respective integration method on two datasets, as they avoided over-fitting, with precision and recall values being more balanced. Regardless of the response, same conclusions were reached, *i*.*e*. implementing multiple sCCA can avoid over-fitting.

Figure 5b presents the scatter-plots of the test set of the canonical vectors of the first canonical variate pair. In contrast with the nutriMouse study, visually there is no clear separation observed between cancer types or survival status of the patients. Since the objective of sCCA is to maximise canonical correlation, it is important to preserve it. The canonical vectors of the test set are computed by linearly combining the estimated canonical vectors (through training), with the original test datasets. RelPMDCCA produced the highest correlation in all three combinations of canonical vectors (Figure 5b).

**Figure 5:**
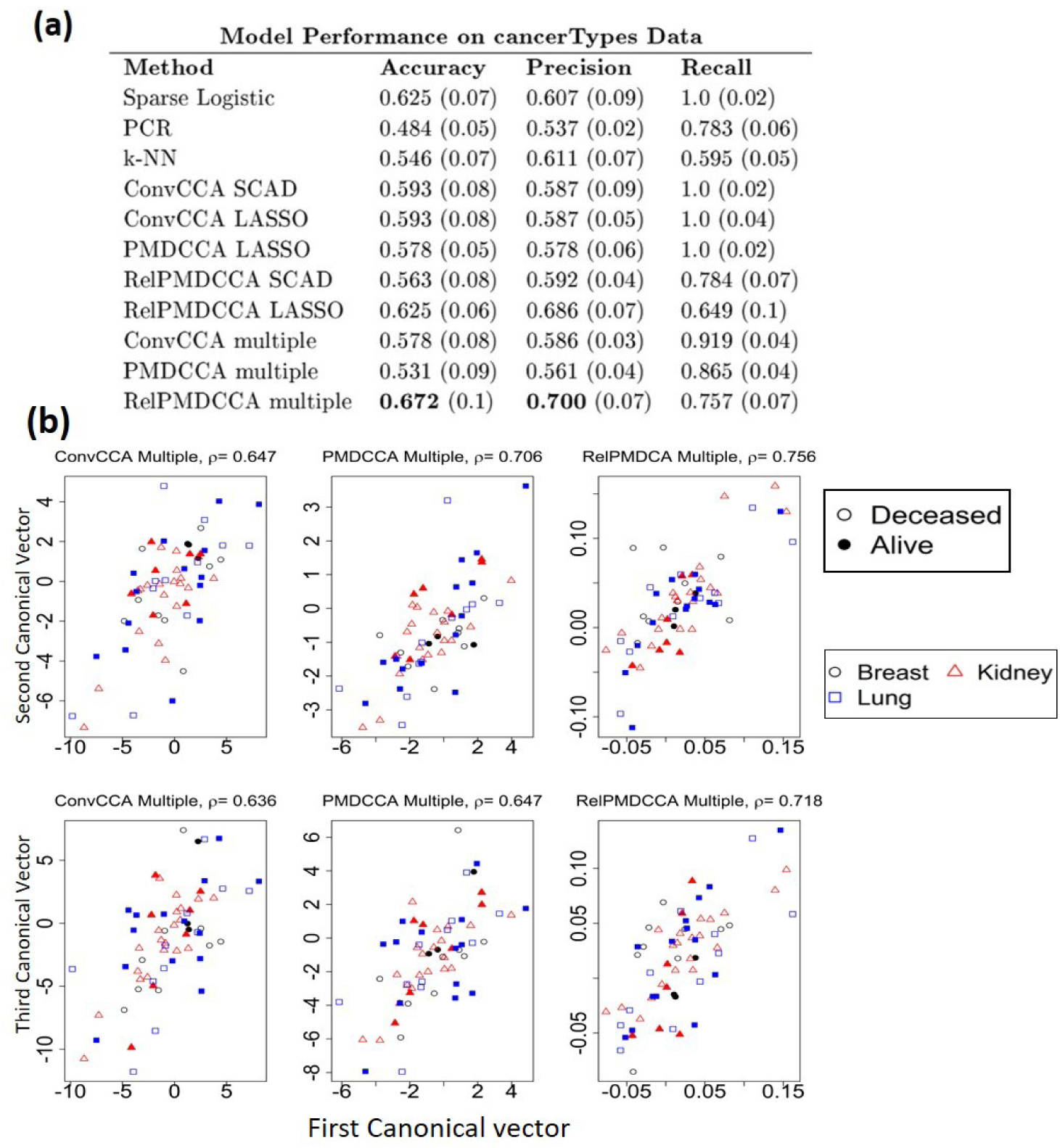
sCCA performance on cancerTypes data. **(a)** Model performance for the prediction of samples’ survival status. The best overall performed model is shown with bold. **(b)** Scatter-plots of canonical variates in cancerTypes analysis through multiple sCCA.

## 5 Discussion

The increasing number of biological, epidemiological and medical studies with multiple datasets on the same samples calls for data integration techniques that can deal with heterogeneity and high dimensional datasets.

In this paper, we investigated and extended three sCCA methods: ConvCCA, PMDCCA and its extension RelPMDCCA. We modified RelPMDCCA to penalise canonical vectors through SCAD and we compared its performance against LASSO penalty. Simulated data using three different models were generated to compare the performance of the three methods under various scenarios. Further, we proposed an extension in computing the additional canonical pairs. The extension satisfies necessary conditions in achieving complete orthogonality among them. Finally, we extended ConvCCA and RelPMDCCA for integrating more than two datasets instead of just two as their original version.

Through both the analysis of simulated and real datasets we have illustrated the benefits of integrating multiple datasets. By collectively reducing the dimensions of the datasets, while obtaining maximal correlation between the datasets, sCCA methods were found to have better accuracy in predicting complex traits than other conventional machine learning methods. Through our proposed extensions of the ConvCCA and RelPMDCCA approaches for integrating more than two datasets and for incorporating the response matrix as one of the integrated datasets, we have showed that over-fitting can be avoided and higher predictive accuracy can be obtained.

Through the analysis of the two real datasets we illustrated that the sCCA methods can improve our predictions of complex traits in both cases: (*i*) when a regression model is built with the new canonical matrices as input matrices and (*ii*) when the response matrix is one of the input matrices in the data integration. For both cases, the sCCA methods can improve the predictions of the response.

Our simulation studies findings are in agreement with Chalise and Fridley [2012] that showed that ConvCCA has better results with SCAD penalty rather than LASSO. When dealing with two independent datasets, ConvCCA may be the favoured method, but with multiple independent datasets PMDCCA showed the most reliable results. With no exception, RelPMDCCA provides the highest canonical correlation in all simulation studies and all real-data analyses performed in this paper. In addition to this observation, RelPMDCCA also provides a sparser outcome.

To preserve orthogonality, the results of our simulation study suggest different methods based on data characteristics. If the data satisfy *n* > *p*_1_ > *p*_2_, then ConvCCA is a more sensible choice. In the other cases (*p*_1_ > *n* > *p*_2_ or *p*_1_ > *p*_2_ > *n*), ConvCCA failed to provide orthogonal canonical pairs, while PMDCCA and RelPMDCCA, attained orthogonality on synthetic data from all three data generating models. We observed the performance of the sCCA methods to depend on the structure of the input data. For sparse datasets we recommend the use of the RelPMDCCA approach as it is the one that performed better for such datasets.

Huang et al. [2017] argued the need for a comparison of data integration approaches. This paper has addressed this by evaluating the performance of three sCCA approaches. We have further illustrated that integrating datasets through multiple sCCA could improve the prediction power, suggesting that researchers with access to two or more datasets should aim for an integrative analysis.

## Acknowledgements

The authors would like to thank Sarah Filippi and Philipp Thomas for useful conversations.

## Supplementary Material

### S1. Selection of tuning parameters

As with all penalisation methods, selecting the tuning parameters is vital in the performance of the algorithm. In sCCA methods, it is necessary to select the optimal choices for the parameters (*τ*_*w*_) of all input data-sets, which are likely to differ from each other and influence one another.

To achieve this, *k*-fold cross-validation is performed on different values for those parameters. The aim of sCCA methods is to maximise the canonical correlation and thus the selection of tuning parameters is based on that. Suppose we have *X*_1_ and *X*_2_, then the following measure is taken for every choice of 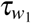 and 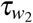, usually within a range of values in (0, 0.3):

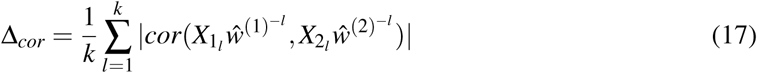

where 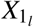 and 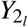 represents the testing sets of *X*_1_ and *X*_2_ for fold *l*, respectively, and 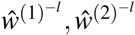 the estimated canonical variate pair based on the training sets.

In order to determine the optimal tuning parameters at each run of the algorithms, one must first compute Δ_*cor*_ for all choices of 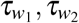 for all *k* folds. The values of 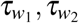 that maximise Δ_*cor*_ are then taken as optimal.

Due to the iterative nature of the algorithms, the choice of 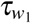 will influence the final outcome of *X*_2_ as well. Hence, selecting the optimal tuning parameters in a multiple-data setting is more complicated and computationally heavy. With two data-sets, 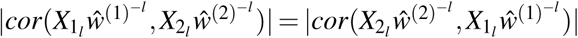, and so we can compute Δ_*cor*_ once for every combination of 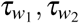 in each fold. With *M* data-sets, Δ_*cor*_ is computed *M* times. In multiple sCCA, eq. 17, is replaced by:

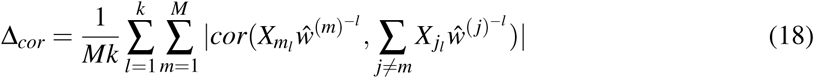

The time complexity of selecting tuning parameter in multiple data-sets is notably high. To reduce it, a threshold in the correlation values was used. Even though the optimal selection may not be guaranteed, well-performed tuning parameters are selected based on the threshold.

### S2. Evaluation measures

The closer the estimated canonical variate pairs are to the true pairs, the better the performance of sCCA methods. We took similar measures as Bonner and Beyene [2017] used in evaluating sparse PCA approaches.

The primary criteria in the evaluation will be the classification of zero-valued and non-zero-valued elements of the canonical vectors, since these would signify the grouping structure. In addition to the structure of the estimated **ŵ**^(1)^ and **ŵ**^(2)^, it is important to estimate values close to the true ones. Hence, we also measured numerical differences between true and estimated values of the canonical vectors.

In particular, we examined the performance of the simulations based on the following measures:

1. **NZ**: The number of non-zero values remaining in the estimated pairs. Expecting a sparse representation
2. **TRUENZ**: The number of correctly classified **non-zero** values
3. **TRUEZ**: The number of correctly classified **zero** values
4. **ANGLE**: A measure of the distance between true and estimated canonical variate pairs. A value between 0 (perfect) and 1(worst) It is calculated for each **w**^(*i*)^, separately as follows: 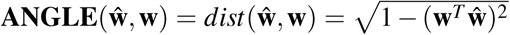, where **w** is the true canonical variate and **ŵ** is the estimated one.
5. **LOSS** (X. Suo *et al*., 2017): A loss value between the true and the estimated canonical correlation pairs. For each element of the pair, **w**, the loss is computed as follows:

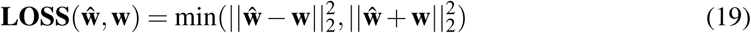
6. **CORR**: The estimated canonical correlation *Cor*(*X*_1_**ŵ**^(1)^, *X*_2_**ŵ**^(2)^) Note that all measures except **CORR**, were computed separately for all **ŵ**^(*i*)^, *i* = 1, 2.

The measures **TRUENZ** and **TRUEZ** are not very intuitive on their own, especially since simulations were conducted with different parameters. That is, a different true number of non-zero values are to be identified, depending on the scenario performed. Hence, a confusion matrix of the zero and non-zero elements found, against the simulated truth, was computed. True Positive Rate (**TPR**), False Positive Rate (**FPR**), Positive Predictive Value (**PPV**), Negative Predictive Value (**NPV**) and Accuracy (**ACC**) were then obtained.

Overall, the evaluation measures cover the entire performance of the methods: the correct identification of non-zero (and zero) values, the exact values of the canonical vectors (through **ANGLE** and **LOSS**) and the correlation within the canonical pairs.

### S3. Supplementary Figures

**Figure 6:**
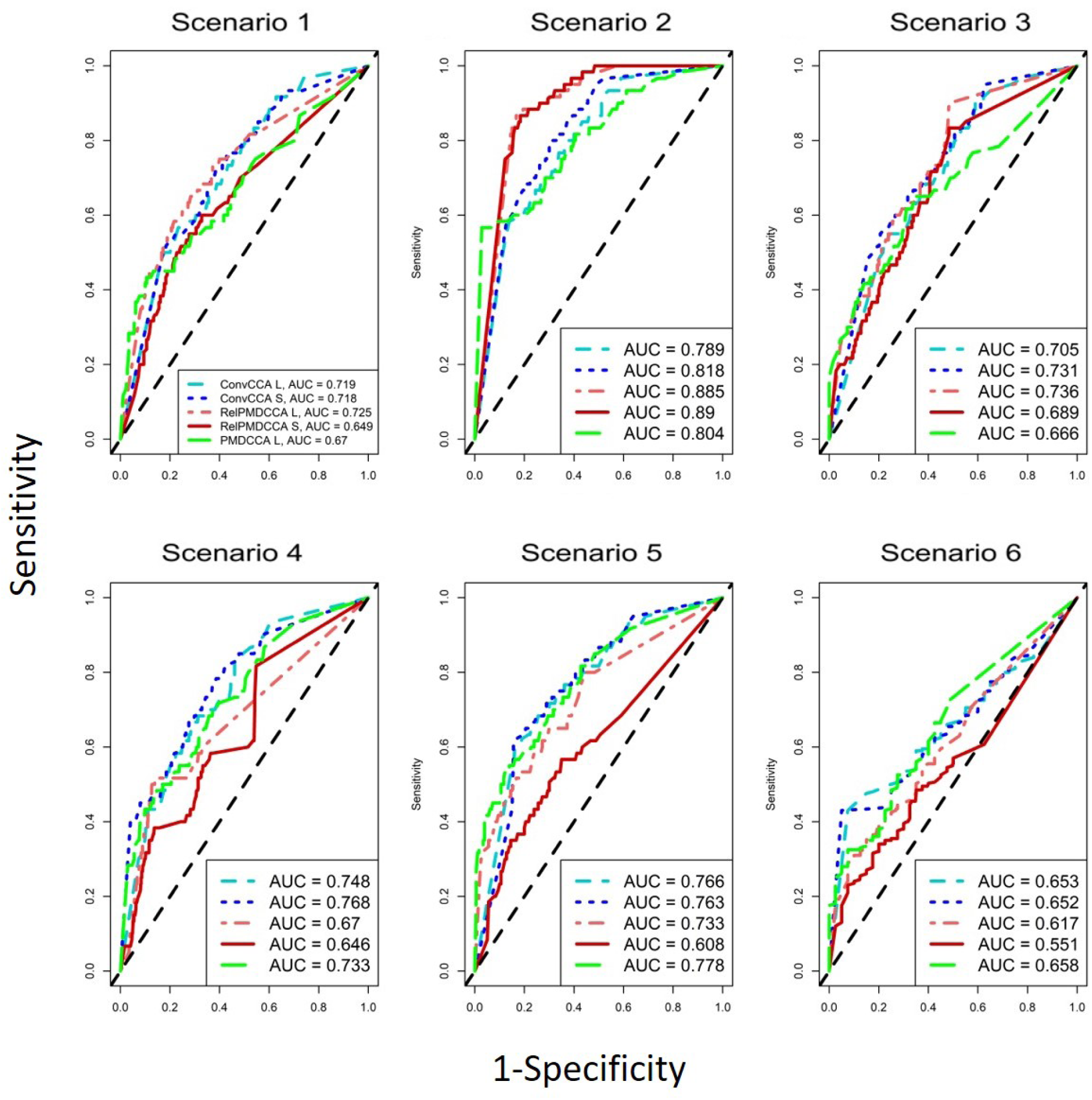
ROC curve plots, showing averaged results (over the models) for each scenario on *X*_2_**w**^(2)^.

**Figure 7:**
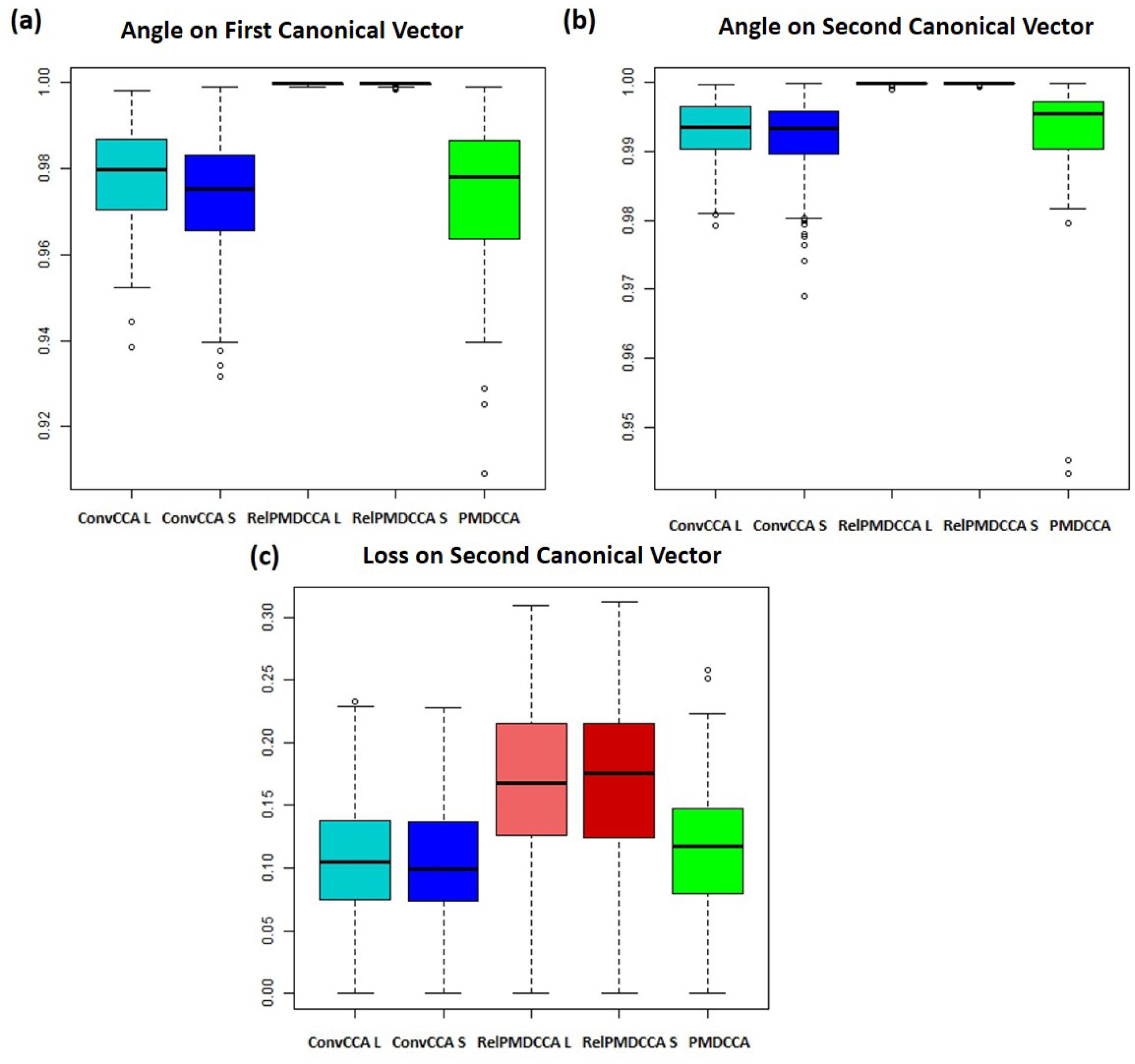
**(a)** Angle evaluation measure on the first canonical vector(*X*_1_**w**^(1)^). **(b)** Angle on the second canonical vector (*X*_2_**w**^(2)^) **(c)** Loss on the second canonical vector

**Figure 8:**
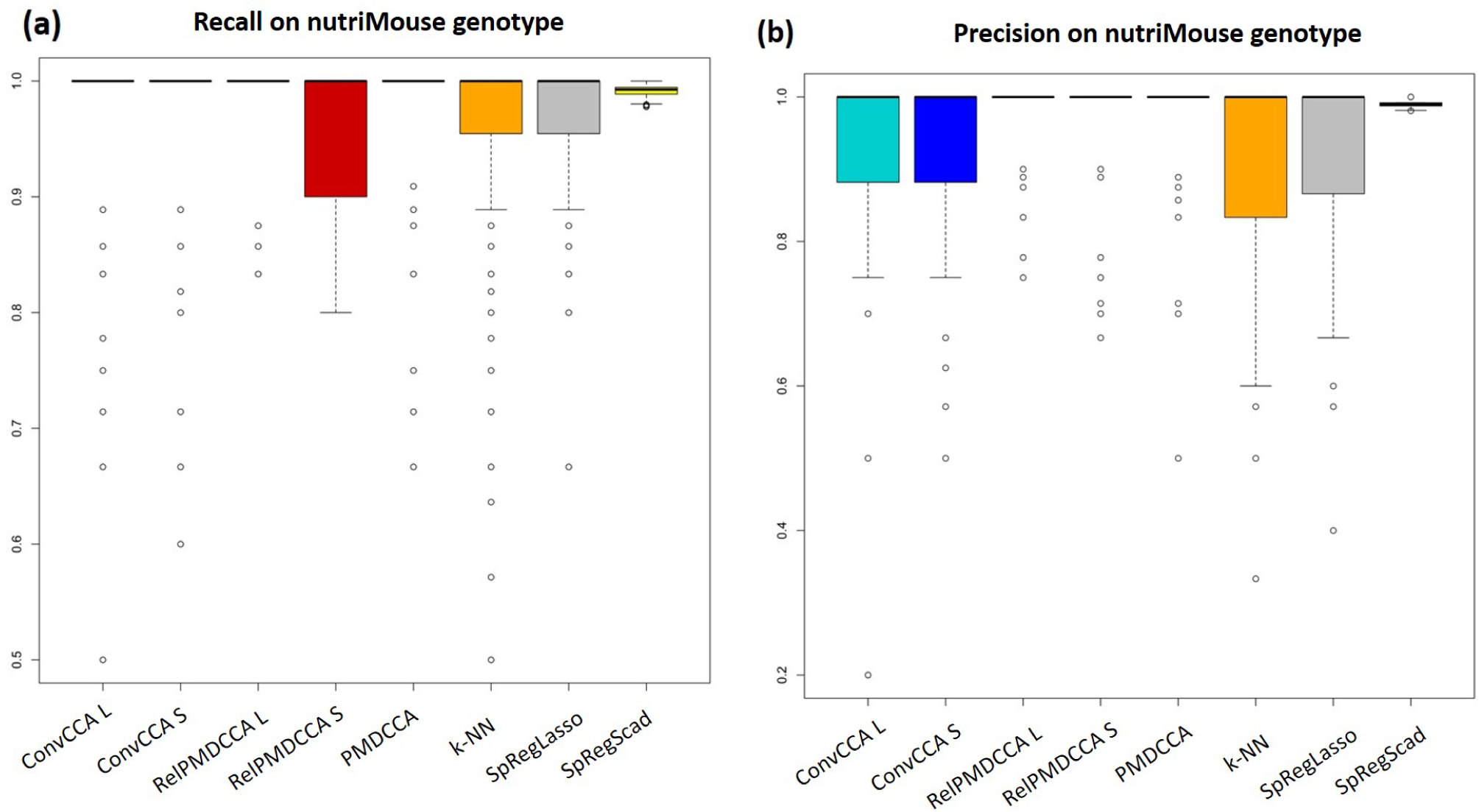
**(a)** Recall and **(b)** precision measures in nutriMouse study in predicting the genotype of mice

**Figure 9:**
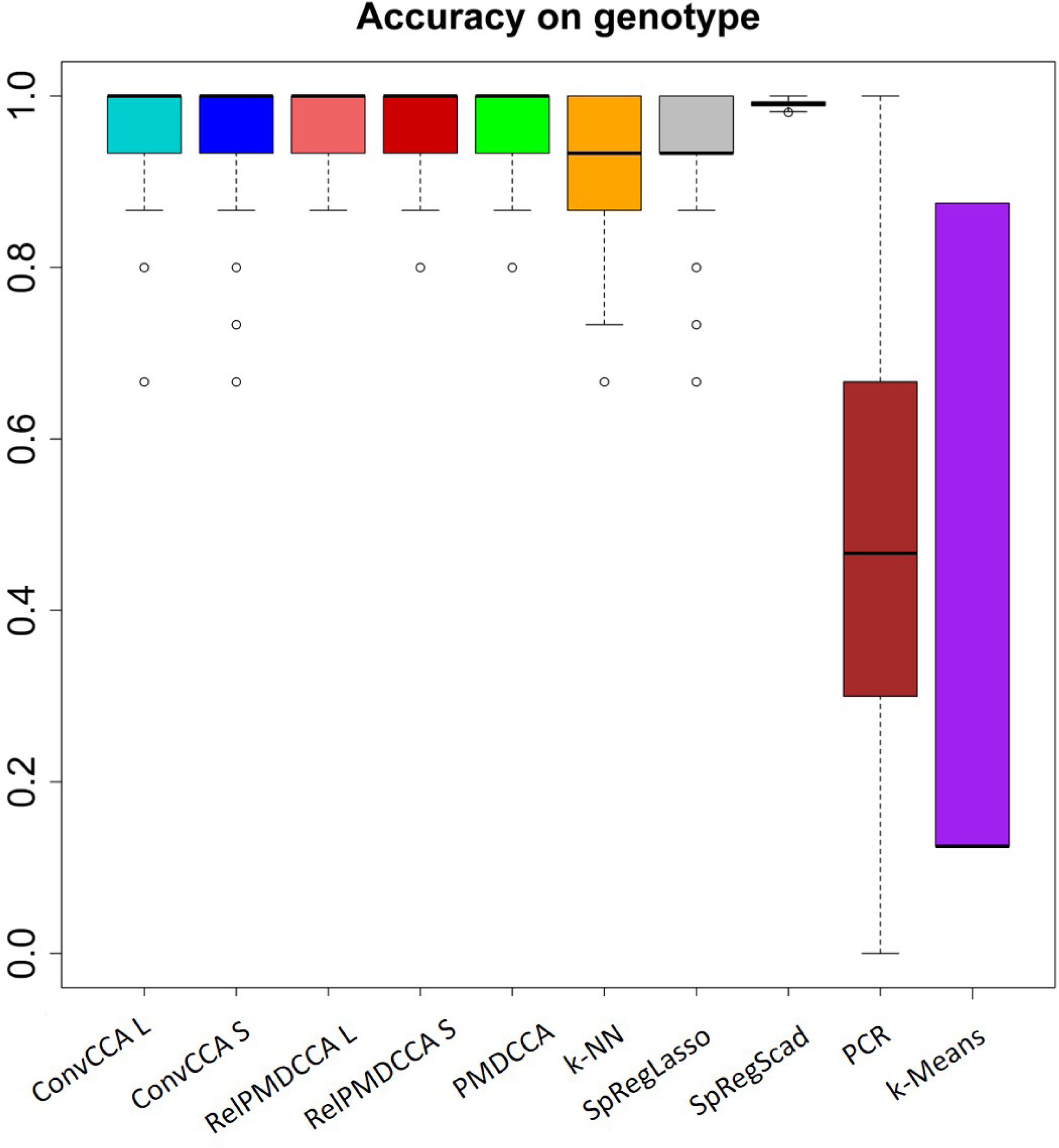
Accuracy of all learning models (including PCR and k-means) in predicting the genotype of mice in the nutriMouse studys

**Figure 10:**
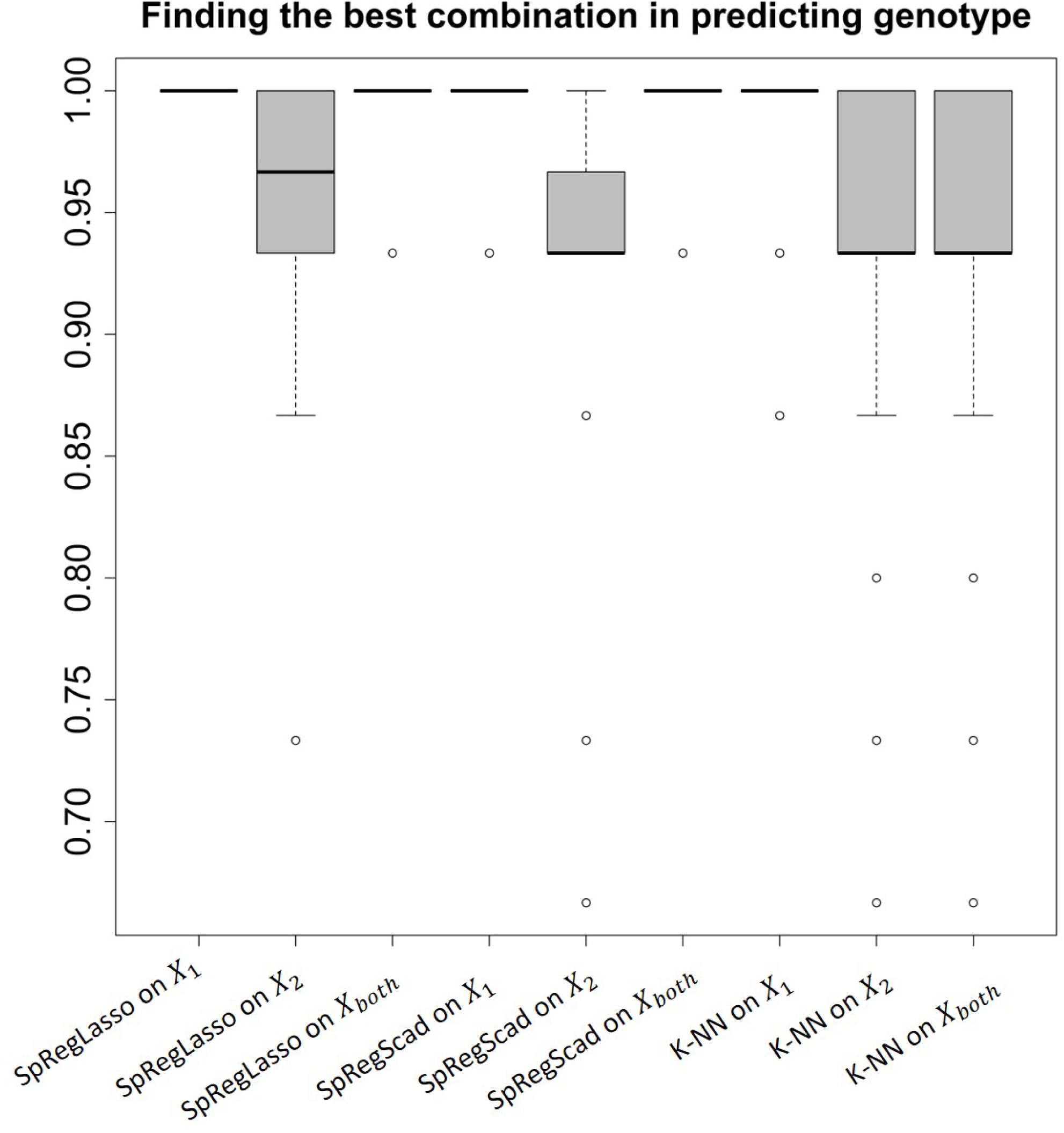
Accuracy measures on SpRegLasso, SpRegScad and k-NN with *X*_1_, *X*_2_ and *X*_*both*_ acting as predictors, with the response being the mice genotype from the nutriMouse study.

**Figure 11:**
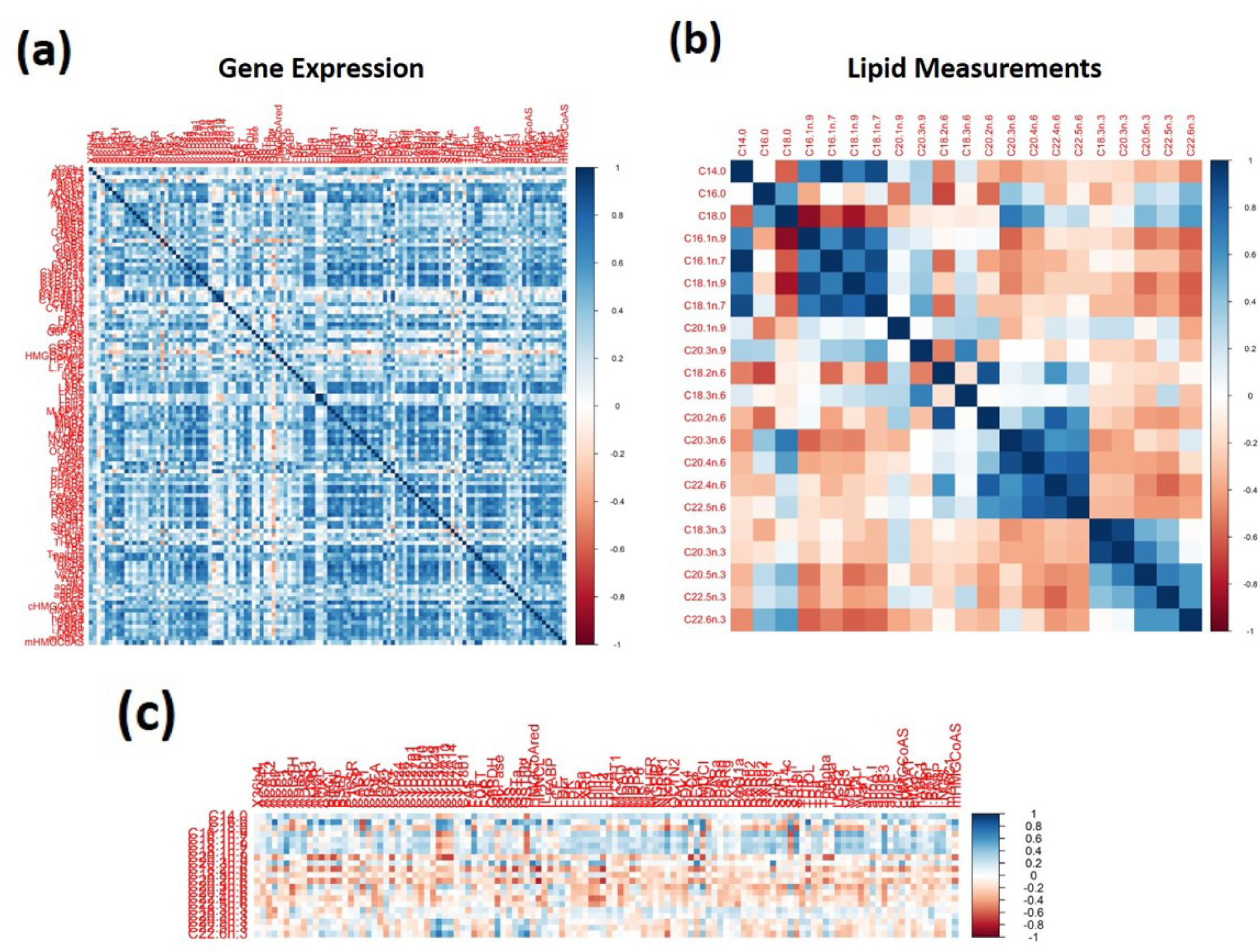
Correlation plots of **(a)** gene expression, and **(b)** lipid measurements in nutriMouse study. **(c)** Cross-correlation of the two data-sets.

